# The striatum drives the ergogenic effects of caffeine

**DOI:** 10.1101/2022.10.06.511163

**Authors:** Ana Cristina de Bem Alves, Ana Elisa Speck, Hémelin Resende Farias, Leo Meira Martins, Naiara Souza dos Santos, Gabriela Pannata, Ana Paula Tavares, Jade de Oliveira, Ângelo R. Tomé, Rodrigo A. Cunha, Aderbal S Aguiar

## Abstract

Caffeine is one of the main ergogenic resources used in exercise and sports. Previously, we reported the ergogenic mechanism of caffeine through neuronal A_2A_R antagonism in the central nervous system [1]. We now demonstrate that the striatum rules the ergogenic effects of caffeine through neuroplasticity changes. Thirty-four Swiss (8-10 weeks, 47 ± 1.5 g) and twenty-four C57BL/6J (8-10 weeks, 23.9 ± 0.4 g) adult male mice were studied behaviorly and electrophysiologically using caffeine and energy metabolism was studied in SH-SY5Y cells. Systemic (15 mg/kg, i.p.) or striatal (bilateral, 15 μg) caffeine was psychostimulant in the open field (*p* < 0.05) and increased grip efficiency (*p* < 0.05). Caffeine also shifted long-term depression (LTD) to potentiation (LTP) in striatal slices and increased the mitochondrial mass (*p* < 0.05) and membrane potential (*p* < 0.05) in SH-SY5Y dopaminergic cells. Our results demonstrate the role of the striatum in the ergogenic effects of caffeine, with changes in neuroplasticity and mitochondrial metabolism.

## INTRODUCTION

Exercise fatigue is one of the main barriers to exercise and sports performance. Caffeine (1,3,7-trimethyl xanthine) is a non-selective antagonist of adenosine A_1_ and A_2A_ receptors (A_1_R and A_2A_R) that exhibit psychostimulant and ergogenic effects (i.e., increases physical performance) [1]. It is one of the most popular ergogenic aids in exercise and sports, available in coffee, tea, caffeinated soft and sports drinks, sports supplements, and other beverages and foods [2–5]The gastrointestinal system rapidly absorbs caffeine, distributed throughout the body fluids, reaching plasma peak concentrations of 2-15 μM (100 mg caffeine) with a half-life of 2.5-10h [6, 7]. The World Anti-Doping Agency (WADA) detected caffeine in 73.8% of 20,686 urine samples from elite athletes between 2004 and 2008 [8], a percentage close to that reported by Van Thuyne et al. [9]: 73.6% in 11,361 samples between 1993 and 2002. The International Society of Sports Nutrition recommends the oral consumption of caffeine in an anhydrous form in doses between 3-6 mg/kg (100-300 mg) [10]. Larger doses (> 9 mg/kg) do not enhance the ergogenic effect and are unsafe as they can trigger adverse effects problematic for athletic performance, such as restlessness, gastrointestinal discomfort, nausea, insomnia, anxiety, and chest pain [10].

Several of the central effects of caffeine can explain the ergogenic effects of caffeine. In fact, caffeine-enhancing effects include improved neurological functions such as typing speed, simple reaction time, sustained attention, memory, and logical reasoning during rest [11]. The physiological targets of caffeine for micromolar concentrations achieved with dietary consumption are A_1_ and A_2A_ adenosinergic receptors [6, 7]. Davis first demonstrated that the adenosine receptor agonist 5⍰-N-methyl carboxamide adenosine (NECA), injected into the rat ventricles, increases running performance [12]. We then demonstrated the role of endogenous extracellular adenosine in the development of fatigue in mice [13] and the pivotal role of neuronal A_2A_R in the mouse forebrain for the ergogenic effects of caffeine [1, 14]. Caffeine increases performance expectancy in cyclists [15], reinforcing the findings of Sun et al. [16] on the role of A_2A_R in the *nucleus accumbens* (*NAc*) on early fatigue associated with exertion in mice or effort-based cost-benefit decision-making (E-CBDM). Accordingly, the overfunction of A_2A_R [17, 18] is a hypothetical mechanism for the fatigue symptom (E-CBDM) in neurological diseases [19].

Fatigue recruits corticostriatal loop connections that reduce performance [20] and impairs long-term potentiation (LTP) and long-term depression (LTD) [21]. This exercise-induced alterations of adenosine signaling [1, 13] and of neuroplasticity [20, 21] are candidate mechanisms responsible for the central component of fatigue. In this work, we investigated the striatal neuroplasticity of caffeine ergogenicity.

## MATERIALS AND METHODS

Four different experiments were conducted in three different laboratories. First, caffeine was administered *in vivo* through systemic (i.p., experiment #1) or local treatment via stereotaxy (striatum, experiment #2) to assess their impact on animals’ fatigue and motor control (Swiss mice, Labioex/UFSC, Brazil). Extracellular electrophysiology analyzed neuroplasticity in striatal slices treated with caffeine (experiment #3, C57BL/6J mice, CNC/U Coimbra, Portugal), and experiment #4 evaluated *in vitro* the effects of caffeine on the mitochondrial metabolism of SH-SY5Y human neuroblastoma cell lines (UFGRS/Brazil).

### Experimental designs #1 and #2

#### Animals, caffeine, and stereotaxy

We used 34 Swiss naïve male adult mice (8-10 weeks, 47 ± 1.5 g). The animal housing had controlled lighting and temperature (12h light-dark cycle, lights on at 7:00 and off at 19:00, and room temperature of 21±1°C) and *ad libitum* access to food and tap water. Housing and handling of the animals followed the current Brazilian legislation (National Council for the Control of Animal Experimentation -CONCEA) and received bioethical approval (CEUA 1503210519). The allocation of animals to the experimental groups was random. Systemic treatment (in a volume of 10 ml/kg) with caffeine (15 mg/kg, i.p.) or saline (NaCl 0.9%, i.p.) was performed 15 min before the behavior and exercise tests.

Another group of mice were anesthetized with ketamine/xylazine, and two cannulas were implanted, one in the right striatum (AP 0.5 mm; ML 2 mm and DV -3 mm) and other in the left striatum (AP 0.5 mm; ML -2 mm and DV -3 mm) [22]. One week after surgery, 4 μL of caffeine (15 μg) or saline (0.5 μg) was injected into conscious animals using an infusion pump (2 μL/minute, Bonther®, Ribeirão Preto, Brazil) immediately before behavioral tests. The caffeine dose (15 μg) was selected through pilot studies and literature data [23, 24]. The mortality rate was 30% (N=9).

#### Behavioral analysis

The behavioral experiments were performed during the light phase of the circadian cycle, after 1 hour of habituation, between 9:00 and 16:00, under controlled temperature (22±1ºC) and luminosity (10 lux). All equipment was cleaned with EtOH 20% after each animal test.

#### Open field

We manually recorded crossings and rearings in the circular open field test (diameter 58 cm × height 50 cm, Insight® EP154C, Ribeirão Preto, Brazil) for 5 minutes [25].

#### Rota rod

The Rotarod test (Insight® EFF411) was used to evaluate motor coordination. The primary inclusion criterion was remaining in the stationary cylinder (0 RPM) for 30 seconds and rotating (5 RPM) for 90 seconds [26]. After 30 minutes, the latency to fall was evaluated in the accelerated cylinder (0.1 RPM/second) with an initial rotation of 5 RPM. Three falls excluded the animal from the evaluation stage. All pieces of equipment were cleaned after each animal test.

#### Grip strength meter

The grip strength meter test (Bonther® 5 kgf) graded the strength and time of the front paw grip. Each mouse was placed individually in the grip meter bar, and after both paws were holding the bar, the experimenter gently pulled the tail in the opposite direction of the bar. We performed 4 trials with 10 seconds each and 1-minute intervals between trials, and we considered the average of 3 best trials (3 out of 4) [27–29]. The equipment measures the force applied and the grip time, and the software provides the maximum and submaximal values. The decrease in physical performance is one of the characteristics of fatigue, evaluated in this behavioral test by the decrease in strength and grip time. We finalized the fatigue assessment by calculating the impulse by integrating the force x grip time curve.

### Experimental design #3

#### Electrophysiology

Twenty-four naïve C57BL/6J mice (male, 23.7⍰± ⍰0.5 g, 8–10 weeks old) were euthanized through cervical dislocation, and the brain was quickly removed (within 1 min) and placed in ice-cold, oxygenated (95% O_2_ and 5% CO_2_) artificial cerebrospinal fluid (ACSF; in mM: 124 NaCl, 4.5 KCl, 1.2 Na_2_HPO_4_, 26 NaHCO_3_, 1.2 CaCl_2_, 1 MgCl_2_, 10 glucose). Striatal slices (400 μm thick) were obtained using a Vibratome 1500 (Leica, Wetzlar, Germany) and allowed to recover for at least 90 min before being transferred to a submerged recording chamber and superfused at 3 mL/min with oxygenated ACSF kept at 30.5ºC. Corticostriatal transmission was assessed with the stimulating electrode placed in the corpus callosum and the recording electrode filled with ACSF (2–5 MΩ) placed in the dorsolateral aspect of the striatum. The neuronal stimulus was delivered with a Grass S44 stimulator (Grass Technologies, RI), and the recordings were obtained by using an ISO-80 amplifier (World Precision Instruments, Hertfordshire, England) and digitized with an ADC-42 system (Pico Technologies, Pelham, NY). Three consecutive population spike responses were averaged and quantified using the Win LTP software v. 2.20b (WinLTP Ltd., Bristol, UK). The relationship between dendritic responsiveness and the synaptic input was determined based on input/output curves in which the population spike amplitude was plotted *versus* the stimulus intensity before and after administering different concentrations of caffeine in the ACSF solution. Stimulation intensity was then selected to yield 40%–50% of the maximum response. Corticostriatal slices were treated with 1 μM, 3 μM or 10 μM caffeine in ACSF oxygenated solution for twenty minutes before the high-frequency stimulation (HFS). HFS consists of three trains each with 100 Hz pulses for 1 s with 10-s intervals between trains. The amplitude of synaptic plasticity was evaluated by the alteration of population spike amplitude 30 min after HFS for each concentration of caffeine [30– 32]. Importantly, the tested concentrations of caffeine only act through the antagonism of adenosine receptors to impact of synaptic transmission [32].

### Experimental design #4

#### Cell culture and evaluation of mitochondrial metabolism

Human neuroblastoma SH-SY5Y cells were maintained in DMEM/F12 supplemented with 10% FBS, 100 units/mL penicillin, 100 μg/mL streptomycin at 37ºC, with 5% CO_2_, under standard conditions. Cells were incubated with PBS or caffeine in concentrations (1 μM, 3 μM or 10 μM) selectively acting as an adenosine receptor antagonist [6, 7]. An MTT assay (see below) evaluated the viability of cells 20 minutes after caffeine exposure [33]. Mitochondrial mass and membrane potential were analyzed using green and red mitotracker [34].

#### MTT assay

SH-SY5Y cells were seeded into 96-well plates in DMEM-F12 with 10% FBS (1⍰× ⍰10_4_ cells/well). After caffeine treatment, the cells were incubated with 0.5 mg/mL of MTT for 2 h at 37 °C. The cell viability was measured by quantifying the activity of cellular dehydrogenases that reduce MTT (3-4,5-dimethyl thiazolyl-2,5-diphenyl-2H-tetrazolium bromide, Sigma Inc.) to a purple formazan salt [35]. The formazan salt formed was dissolved in dimethyl sulfoxide (DMSO), and the absorbance was quantified in a spectrophotometer (Spectra Max 190, Molecular Devices, Sunnyvale, CA, USA) at 540 nm [34].

#### Mitotracker Flow Cytometry Protocol

Mitotracker Green FM (Invitrogen) (MTG) or Mitotracker Red (MTR) are used for staining mitochondria of live cells depending, respectively, on the organelle lipid content (mitochondrial mass) and oxidative activity (mitochondrial membrane potential). Briefly, after caffeine treatment, SH-SY5Y cells were washed on PBS saline and harvested using trypsin. Then, cells were resuspended and incubated for 20 min in the dark with 100 nM of MTG and MTR diluted into pre-warmed (at 37 °C) DMEM F-12. Emitted fluorescence was measured (FL1 ‘green’ 530 nm/30; FL3 ‘red’ 670 nm long pass) in a FACSCalibur using CellQuest Pro software (Becton Dickinson, Franklin Lakes, NJ, USA) and data from 10.000 events were acquired using a log scale for all parameters. The fluorescence mean intensity of FL1-H and FL3-H was evaluated to estimate mitochondrial mass and potential, respectively. All flow cytometry analyses were performed using the FCS Express 4 software (De Novo, Pasadena, CA, USA) [34].

### Statistical Analyses

Statistical analysis was performed using GraphPad Prism version 6.0 for Windows (GraphPad Software, San Diego, California, USA, www.graphpad.com). A one-tailed Student’s t-test analyzed the open field test, rotarod, and grip strength meter; the Cohen’s effect size (*d*) was calculated as small (0.2), medium (0.5), large (0.8), and very large (1.3) [36]. A one-way ANOVA analyzed electrophysiological and mitochondrial assays; Cohen’s effect size (η^2^) was calculated as small (0.01), medium (0.09), and large (0.25) (1.3) [37]. Differences were considered significant when *p*<0.05.

### Data Availability

This work makes open data available under the creative commons license (CC BY) [38].

## RESULTS

### Systemic and striatal treatment with caffeine is ergogenic

Systemic caffeine increased crossings (t_22_= 2.16, *p*< 0.05, *d* = 1.0; Fig.1A) and rearings (t_22_= 2.84, *p*< 0.05, *d* = 1.2; Fig.1B) in the open field, and motor coordination in the rotarod (t_28_= 3.08, *p*< 0.05, *d* = 0.45; Fig.1C). Caffeine did not alter grip strength peak (t_22_= 1.37, *p*= 0.09; Fig.1D) but increased grip time (t_22_= 3.31, *p*< 0.05, *d*= 1.13; Fig.1E). Figure 1F’ shows the Area Under Curve (AUC) from strength *vs*. time results to demonstrate that ergogenic effects of caffeine are due to impulse magnitude (t_22_= 2.14, *p*< 0.05, *d* = 0.81; Fig.1F-F’).

**Fig 1.**
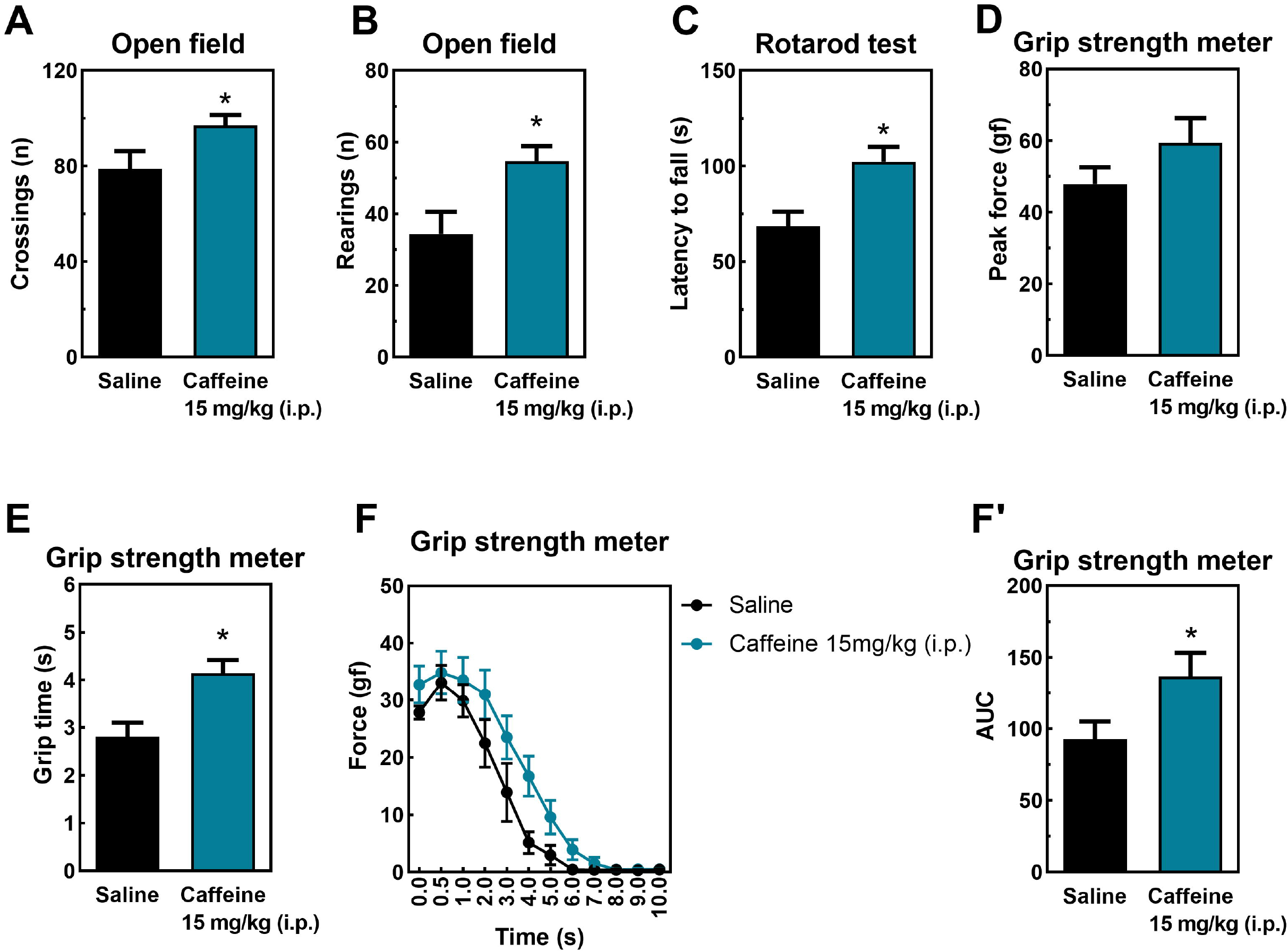
The behavioral effects of systemic caffeine in mice, namely psychostimulating (A and B), improved motor coordination (C), no effect on grip strength (D), but increase in grip time (F). Panel F’ is the integral of strength × time (F), representing increased caffeine-induced impulse. Data are mean ± SEM. N=7-8 animals/group for three independent experiments. *P<0.05 *vs*. saline (Student’s t-test).

Figure 2A (top) shows the coordinates of the mice striatum according to Paxinos and Franklin [39], and Fig.2A (bottom) shows a coronal brain view of a mouse brain, confirming that cannulas were implanted in the expected localization, as previously done [40, 41]. Caffeine injected directly into the striatum enhanced the number of crossings (t_8_= 9.81, *p*< 0.05, *d* = 1.82; Fig. 2B) and rearings (t_8_= 3.66, *p*< 0.05, *d* = 1.50; Fig. 2C) in the open field. Similarly, as occurred for its systemic treatment, the direct intra-striatal caffeine administration did not alter peak strength (t_8_= 0.85, *p*= 0.20; Fig. 2D) but improved time grip time (t_8_= 4.08, *p*< 0.05, *d* = 1.56; Fig. 2E) in the grip strength meter test. The ergogenic effect on impulse magnitude (Fig.2F) was also observed after the intra-striatal direct administration of caffeine (t_8_= 3.50, *p*< 0.05, *d* = 1.48; Fig. 2F’).

**Fig 2.**
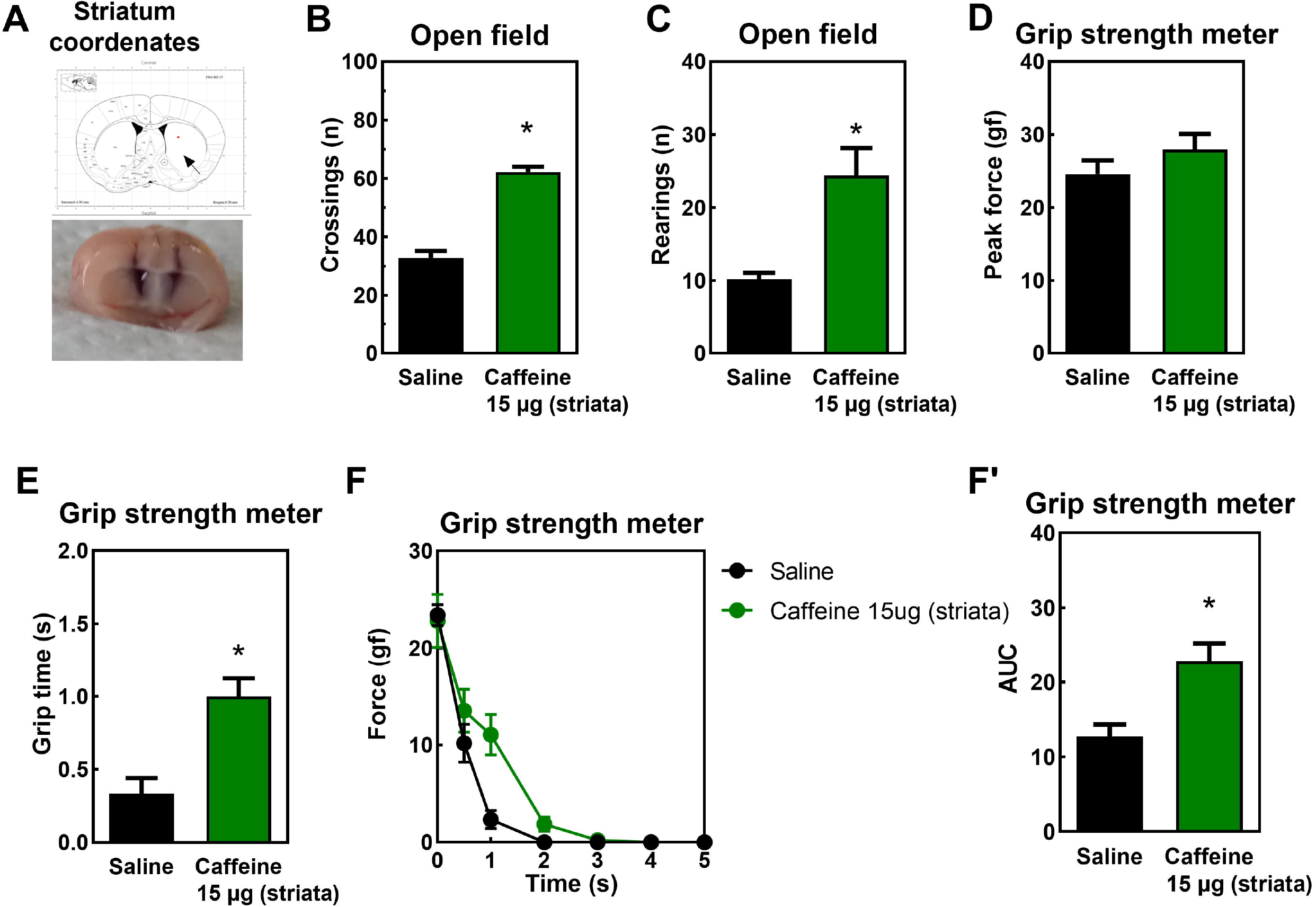
The behavioral effects of intrastriatal (A) injection of caffeine in mice, namely psychostimulating (B and C), no effect on grip strength (D), but an increase in grip time (E). Panel F’ is the integral of strength × time (F), representing increased caffeine-induced impulse. Data are mean ± SEM. N=7-8 animals/group for three independent experiments. *P<0.05 *vs*. saline (Student’s t-test).

### Caffeine changes LTD to LTP in striatum slices

Fig.3A shows the location of the stimulus electrode (top) in the corpus callosum and of the recording electrode (bottom) in the dorsolateral region of the striatum in a coronal brain slice. The first input/output curve (Fig.3B) shows that all slices have a similar pattern of induced excitability during the pre-treatment period without caffeine. After 20 min of caffeine treatment, none of the tested concentrations of caffeine (1-10 μM) significantly altered basal synaptic transmission of corticostriatal synapses (F_3,16_=0.47, p=0.7, Fig.3C). The second input/output curve (Fig.3.D) performed after 20 min of exposure to caffeine shows an increased excitability with the higher concentration of caffeine (10 μM), as would be expected based on the ability of caffeine to antagonize A_1_ receptors controlling basal synaptic transmission [32]. High-frequency stimulation (HFS, black arrow, F_29,864_ =4.26, p< 0.05, η_2_=0.21, Fig.3E) triggered an LTD in control slices (without any treatment), as well as in slices treated with 3 and 10 μM caffeine, whereas the exposure to 1 μM caffeine changed the response to HFS from an LTD to an LTP (F_3.20_ = 11.2, p< 0.05, η^2^=0.62, Fig.3F).

**Fig 3.**
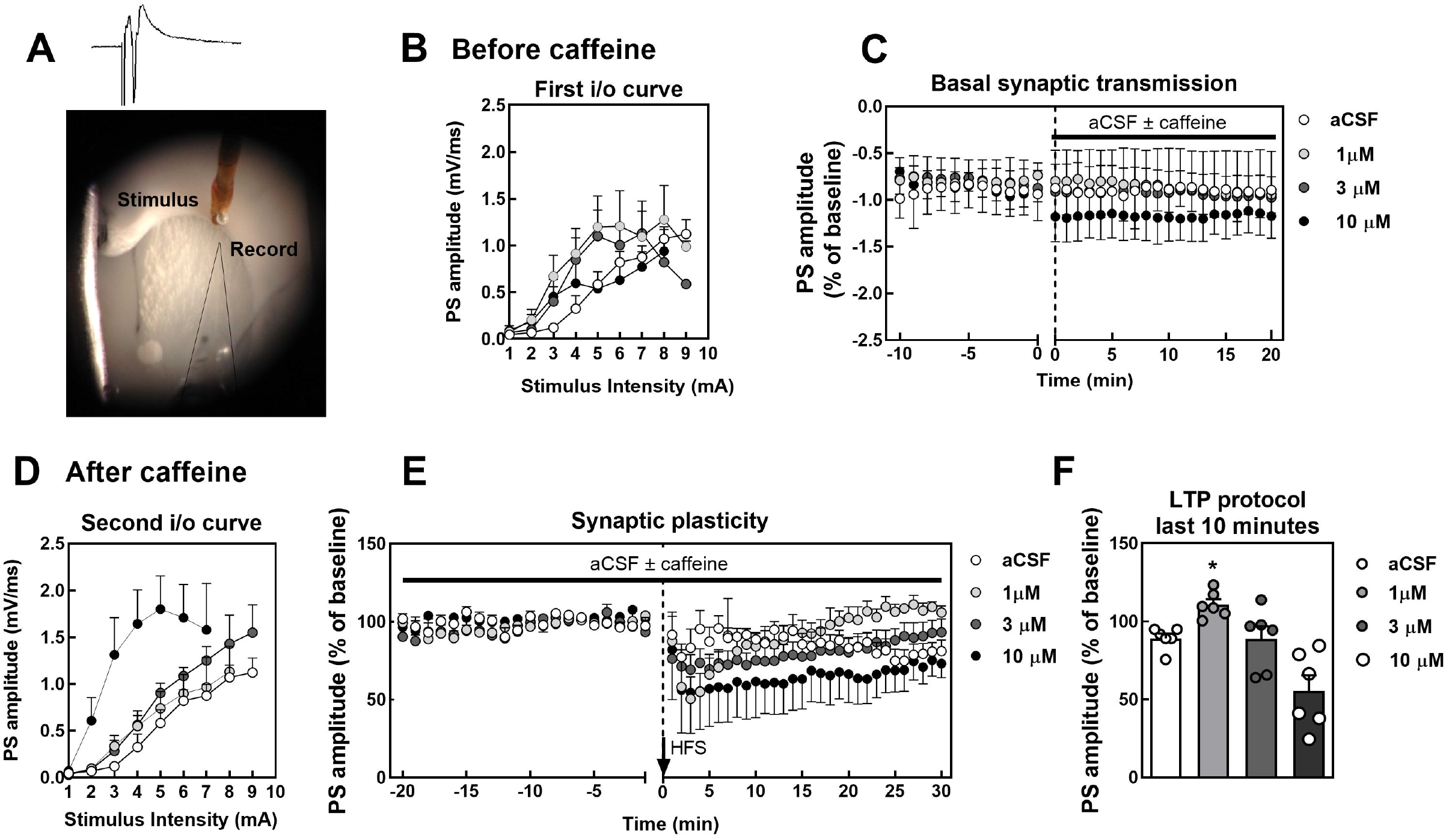
Extracellular electrophysiological recordings of population spike responses at corticostriatal synapses in the dorsolateral striatum in brain coronal slices (A). All groups displayed superimposable input/output curves, indicating similar efficiency of corticostriatal transmission (B). Alteration of basal synaptic transmission after exposure to different concentrations of caffeine (C). Twenty minutes after caffeine infusion, only 10 μM caffeine increased synaptic efficiency (D). The induction of synaptic plasticity by high-frequency stimulation (HFS) caused a long-term depression (LTD) in corticostriatal synapses (E), which was shifted into a long-term potentiation (LTP) upon exposure to 1 μM caffeine, as illustrated in the time course (E) and in the average alterations of the amplitude of synaptic plasticity 30-40 min after HFS (F). Data are means ± SEM (n=6 slices/group). One-way ANOVA followed by Newman–Keuls post hoc tests). aCSF – artificial cerebrospinal fluid. i/o – input and output. PS – pop spike population.

### Caffeine improved mitochondrial metabolism in SH-SY5Y cells

Caffeine acutely increased the mass and membrane potential of the mitochondria of SH-SY5Y cells. Cell viability was not modified after exposure to any tested concentration of caffeine (1 μM, 3 μM or 10 μM), as analyzed by the MTT assay (F_3,68_ = 0.58, *p*= 0.6, η_2_ = 0.03; Fig.4A). Caffeine (1 μM) increased mitochondrial membrane potential (F_3,10_ = 14.99, *p*< 0.05, η_2_ = 0.83; Fig.4B) and mitochondrial mass (F _3,11_= 20.87, *p*< 0.05; Fig.4C) of SH-SY5Y cells. Fig.4F shows a fluorescence shift to the right upon exposure to 1 μM caffeine.

**Fig 4.**
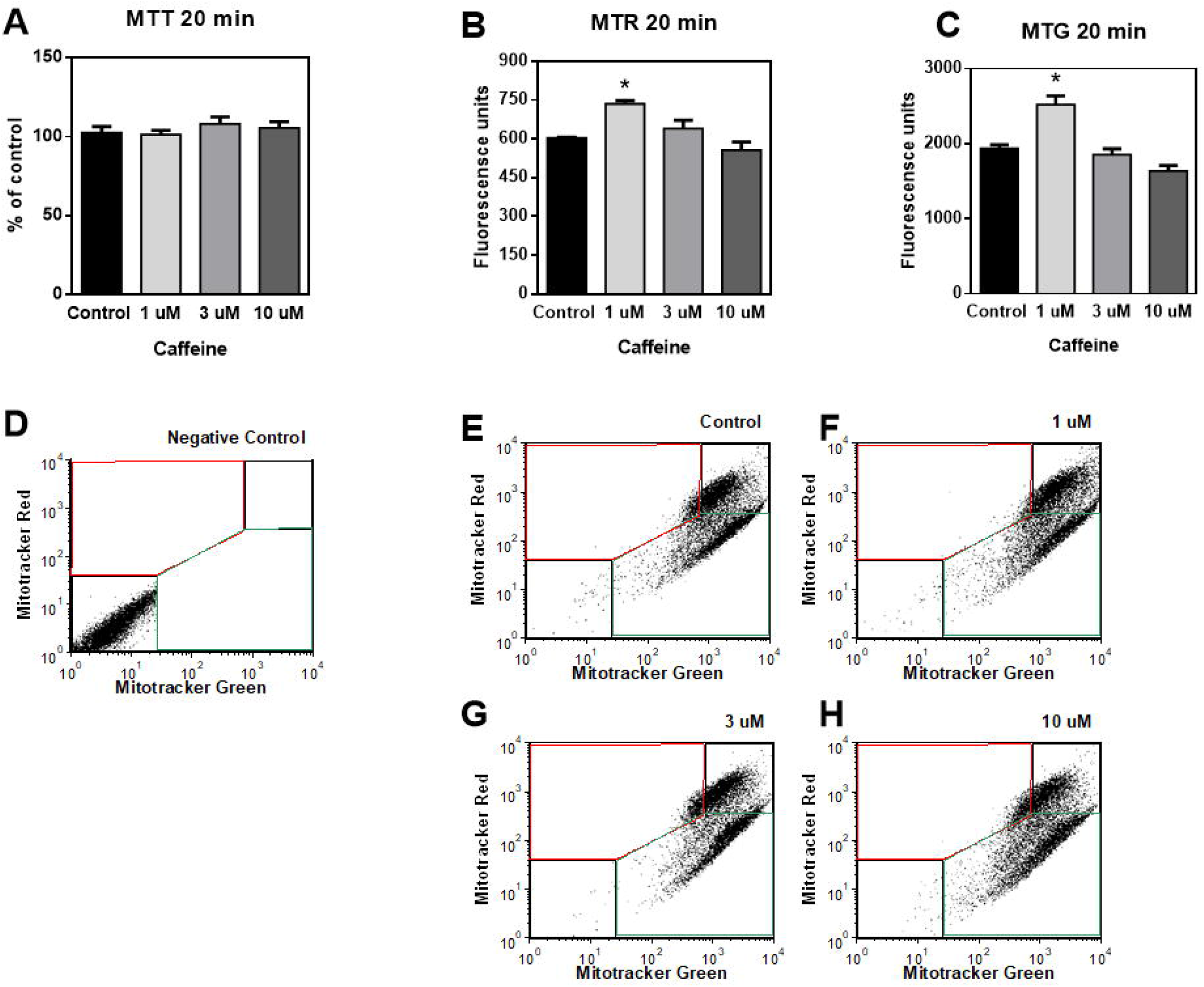
Caffeine improved mitochondrial metabolism in SH-SY5Y cells. Whereas it did not change cell viability (A), caffeine (1 μM) increased mitochondrial membrane potential (B) and mitochondrial mass (C). Panels D-H represents the flow cytometry analyses. Data are means ± SEM. One-way ANOVA followed by Newman–Keuls post hoc tests). MTG - Mitotracker Green. MTR - Mitotracker Red. MTT (3-(4,5-dimethylthiazol-2-yl)-2,5-diphenyltetrazolium bromide).

## DISCUSSION

This work demonstrates that the ergogenic effect of acute systemic and striatal caffeine on grip strength is associated with a change LTD to LTP in the striatum, probably with an increase of the mass and membrane potential of mitochondrial in neurons. Our behavioral data reinforce the evidence that systemic caffeine administration is ergogenic [2–5, 12, 25]. However, we emphasize that this previous knowledge analyzed treadmill running performance, and we demonstrate here for the first time that caffeine increases (grip) muscle strength in mice, an effect also observed in humans [42, 43].

Additionally, we demonstrated for the first time that direct caffeine administration in the striatum reproduces the ergogenic effects upon systemic treatment with caffeine. Statistical effect sizes were large (systemic) and very large (intrastriatal) for caffeine ergogenicity. The modification of force performance was on impulse the grip strength size did not change. Once again, we demonstrate that caffeine modifies a time-derived index of physical performance (force × time), as we have previously demonstrated with running power (force ÷ time) [25].

Overall, we have been building a body of evidence to unravel the neurological mechanisms of the ergogenic effects of caffeine. The ergogenic effects of caffeine depend on the antagonism of neuronal A_2A_R in the rodent CNS [1, 12], and we now show that they involve a local neuroplasticity effect in the striatum, a brain region dense in A_2A_R [44]. Thus, caffeine improves motor coordination (in the rota rod) and grip time while shifting striatal neuroplasticity from an LTD to a LTP. This joins previous findings that caffeine injected into rat brain ventricles improves treadmill performance, an effect that disappears upon administration of the adenosine receptor agonist NECA [12]. Our results correspond to acute effects of caffeine in naïve animals; importantly, previous evidence suggests additional benefits when caffeine is consumed chronically, namely an increased resistance to stress [45, 46], from social [47] to cold stress [48]. In humans, the ergogenic effects of caffeine do not disappear with chronic consumption [2–5]. However, caffeine might increase anxiety [47]which is associated with impaired exercise performance due to reduced processing efficiency, including attention, and increased physiological cost [49, 50]. However, caffeine does not impair exercise performance; on the contrary, it enhances it. The key to understand this apparent paradox might be the arousal effect of caffeine [51]. Physiological arousal in exercise bolsters performance when limited by awareness of exertion [49, 50]. Accordingly, caffeine decreases the rating of perceived exertion [52–54].

Our results strengthen the hypothesis about the ergogenic mechanism of caffeine involve an antagonism of the adenosine modulation system in the central nervous system [12–14, 25]. Central A_2A_R are linked to behavioral responses to exertion [16, 55, 56], such as fatigue and decision making, while A_2A_R antagonists such as caffeine and SCH58261 increase running performance [12, 25], and caffeine-treated forebrain A_2A_R knockouts mice did not demonstrate this ergogenic effect [25]. Indeed, exertion fatigue and ergogenic effects of caffeine are associated with A_2A_R [16, 25, 55, 56]. As the striatum displays a particularly high density of A_2A_R [44], we injected caffeine directly into the dorsolateral striatum of mice, which triggered psychomotor and ergogenic effects. The psychomotor effects of caffeine in the striatum are associated with allosteric modulation of A_2A_R-D_2_R heteromers and presynaptic control of glutamatergic neurotransmission [57]. Striatal A_2A_R facilitate neuronal activity in the striatum by increasing dopaminergic signaling, more by controlling the availability of D_2_ and D_3_ receptors than by influencing dopamine release [58]. Thus, the A_2A_R-D_2_R allosteric interaction modulates psychostimulant responses, which we believe to be involved in the ergogenic effect of caffeine, as discussed above.

Regarding electrophysiological findings, repetition of exercise can modify corticostriatal neuroplasticity, where LTD predominates, which plays an essential role in motor control [59, 60]. Ma et al. [21] demonstrated that exercise-induced fatigue impairs corticostriatal _N_-methyl-_D_-aspartate (NMDA) receptor-dependent LTP and endocannabinoid-dependent LTD (eCB-LTD). We observed that caffeine (1.0 μM) modified *ex vivo* corticostriatal neuroplasticity from LTD to LTP, although this *in vivo* effect remains to be confirmed. Conversely, the increased mitochondrial metabolism observed in cells supports increased neuronal activity at the site, as previously demonstrated in APPsw transgenic mice [61] and neuroblastoma cells [62].

This work is a proof of concept developed in naïve animals that reinforces our working hypothesis that A_2A_R in the corticostriatal pathway play a key ergogenic role [1, 13, 14]. However, some limitations of our work need to be considered. The experiments were carried out in males, a limitation since there is evidence demonstrating that the ergogenic effects of caffeine and A_2A_R antagonists are similar [1, 64–67] or lesser in females [67–69], while the adverse effects are more significant[70]. Moreover, future investigations should evaluate the tolerability developed by regular consumption of coffee and caffeine on ergogenicity. Finally, it should be noted that the present evidence was gathered using different models, whereby behavior was assessed in Swiss mice, slice electrophysiology in C57BL/6J mice and metabolism in human neuroblastoma SH-SY5Y cells. The few studies on the ergogenicity of caffeine do not yet allow us to conclude on the eventual existence of differences between mouse strains, whereas the use of cell culture is expected to allow unveiling metabolic determinants of brain metabolism underlying the ergogenic effect of caffeine.

In summary, our results demonstrate that striatal A_2A_R contribute to the neurophysiology of the ergogenic effects of caffeine.

## DECLARATIONS

### Ethical Approval

The ethics committees approved animal experiments in Brazil (Federal University of Santa Catarina, CEUA 1503210519) and Portugal (University of Coimbra, ORBEA 138-2016/1507201) according to specific legislation.

### Authors’ contributions

ASAJr, JO, and RAC designed the experiment and provided funding. The experiments were performed by ACABA, AES, HRF, NSS, GP, APT, LMM, and ART. ACBA, AES, JO, ART, and AJASr performed the statistics. ACBA, AES, HRF, ART, and ASAJr built the graphics. ACBA, AES, HRF, and ASAJr wrote the manuscript. All authors reviewed the manuscript.

### Availability of data and materials

This work makes open data available under the creative commons license (CC BY) [38].

